# The proto CpG island methylator phenotype of sessile serrated adenomas/polyps

**DOI:** 10.1101/357178

**Authors:** Hannah R. Parker, Stephany Orjuela, Andreia Martinho Oliveira, Fabrizio Cereatti, Matthias Sauter, Henriette Heinrich, Giulia Tanzi, Achim Weber, Paul Komminoth, Stephan Vavricka, Luca Albanese, Federico Buffoli, Mark D. Robinson, Giancarlo Marra

## Abstract

Sessile serrated adenomas/polyps (SSA/Ps) are the putative precursors of the ˜20% of colon cancers with the CpG island methylator phenotype (CIMP), but their molecular features are poorly understood. We used high-throughput analysis of DNA methylation and gene expression to investigate the epigenetic phenotype of SSA/Ps. Fresh-tissue samples of 17 SSA/Ps and (for comparison purposes) 15 conventional adenomas (cADNs)—each with a matched sample of normal mucosa— were prospectively collected during colonoscopy (total no. samples analyzed: 64). DNA and RNA were extracted from each sample. DNA was subjected to bisulfite next-generation sequencing to assess methylation levels at ˜2.7 million CpG sites located predominantly in gene regulatory regions and spanning 80.5Mb (˜2.5% of the genome); RNA was sequenced to define the samples’ transcriptomes. An independent series of 61 archival lesions was used for targeted verification of DNA methylation findings. Compared with normal mucosa samples, SSA/Ps and cADNs exhibited markedly remodeled methylomes. In cADNs, *hypo*methylated regions were far more numerous (18,417 vs 4288 in SSA/Ps) and rarely affected CpG islands/shores. SSA/Ps seemed to have escaped this wave of demethylation. Cytosine *hyper*methylation in SSA/Ps was more pervasive (hypermethylated regions: 22,147 vs 15,965 in cADNs; hypermethylated genes: 4938 vs 3443 in cADNs) and more extensive (region for region), and it occurred mainly within CpG islands and shores. Given its resemblance to the CIMP typical of SSA/Ps’ putative descendant colon cancers, we refer to the SSA/P methylation phenotype as *proto-CIMP*. Verification studies of six hypermethylated regions (3 SSA/P-specific and 3 common) demonstrated the high potential of DNA methylation markers for predicting the diagnosis of SSA/Ps and cADNs. Surprisingly, proto-CIMP in SSA/Ps was associated with upregulated gene expression (n=618 genes vs 349 that were downregulated); downregulation was more common in cADNs (n=712 vs 516 upregulated genes). The epigenetic landscape of SSA/Ps differs markedly from that of cADNs. These differences are a potentially rich source of novel tissue-based and noninvasive biomarkers that can add precision to the clinical management of the two most frequent colon-cancer precursors.

## Introduction

The serrated pathway of tumorigenesis appears to give rise to approximately one out of five sporadic colon cancers—more specifically, those with the CpG island methylator phenotype (CIMP) (Bettington, Walker et al., 2013, Hinoue, Weisenberger et al., 2012, Ijspeert, Vermeulen et al., 2015, Jass, 2005, Okugawa, Grady et al., 2015, Phipps, Limburg et al., 2015). First described in 1999 by Toyota et al.,(Toyota, Ahuja et al., 1999) the CIMP hallmark is a remarkably high level of age-independent cytosine methylation involving CpG dinucleotide clusters (i.e., CpG islands), which are generally unmethylated in normal somatic cells (Deaton & Bird, 2011). Abnormal methylation of these residues is one of the epigenetic changes associated with gene silencing and it has been reported in virtually all types of human cancer (Feinberg, Ohlsson et al., 2006, Issa, 2014, Jones & Baylin, 2007, Rodriguez-Paredes & Esteller, 2011). In CIMP colon cancers, however, this aberrant methylation occurs much more frequently (Hinoue et al., 2012) and is therefore more likely to involve CpG islands within the promoters of genes with functional relevance for tumorigenesis. The DNA mismatch repair (MMR) gene *MLH1,* for instance, is silenced in ˜55% of CIMP colon cancers (Phipps et al., 2015), and its loss results in a hypermutated, microsatellite-unstable phenotype (Bettington, Walker et al., 2017, Boland & Goel, 2010, O’Brien, Yang et al., 2006, Weisenberger, Siegmund et al., 2006) that impacts both the prognosis of the disease and its sensitivity to chemotherapeutics (Carethers, Smith et al., 2004, Le, Uram et al., 2015, Phipps et al., 2015).

At the precancerous level, the World Health Organization (WHO) currently recognizes three classes of serrated colorectal lesions: hyperplastic polyps, traditional serrated adenomas (TSAs), and sessile serrated adenomas/polyps (SSA/Ps) (Bosman, Carneiro et al., 2010). The latter two are both thought to have malignant potential, but SSA/Ps are considered the most likely precursors of CIMP colon malignancies. Like CIMP cancers, large SSA/Ps develop mainly in the proximal colon and frequently express the constitutively activated BRAFV600E serine/threonine kinase (Bettington et al., 2013, Kambara, Simms et al., 2004, Rex, Ahnen et al., 2012, Weisenberger et al., 2006). The hyperactive MAPK signaling caused by this mutant protein is thought to be responsible for many features of the serrated colorectal tumors, including the CIMP (Carragher, Snell et al., 2010, Fang, Ou et al., 2014, Rad, Cadinanos et al., 2013, Sakamoto, Feng et al., 2017).

Early endoscopic removal of advanced precancerous lesions has proved to be the most effective way to reduce colon cancer-related mortality, and large proximal-colon SSA/Ps are now considered no less dangerous in this sense than cADNs, which are the most frequent colon cancer precursors (Lieberman, Rex et al., 2012). However, SSA/Ps are easier to miss during endoscopy, partly because of their location but also because of their morphology (East, Vieth et al., 2015, Gourevitch, Rose et al., 2018, Rex et al., 2012, Singh, Nugent et al., 2010). They are nearly always flat or sessile lesions with poorly defined borders and colors resembling those of the normal mucosa, features that impede both their detection and complete endoscopic resection (Pohl, Srivastava et al., 2013). Finally, pathologic diagnosis of serrated lesions is still subject to substantial inter-examiner variability, so endoscopically resected SSA/Ps are likely to be misclassified (Bettington et al., 2013, Bettington, Walker et al., 2014, Gourevitch et al., 2018, Rex et al., 2012). These factors converge to create an obvious gap in colorectal cancer prevention, reflected by the significant over-representation of the CIMP and proximal-colon locations among interval cancers (i.e., those diagnosed within 2 years of a negative colonoscopy) (Sawhney, Farrar et al., 2006).

Closure of this gap could be hastened by the identification of robust sets of molecular SSA/P biomarkers: stool-based diagnostic panels for use in noninvasive pre-colonoscopy screening programs, and tissue-based panels for more reliable diagnosis of these lesions in pathology laboratories. DNA methylation changes offer several advantages for use as pre-colonoscopy screening markers. Not only are they the most stable of known epigenetic marks, (Bird, 1978),(Wu & Zhang, 2014) they also occur early in tumorigenesis (Feinberg et al., 2006, Issa, 2014, Jones & Baylin, 2007, Menigatti, Cattaneo et al., 2009, Menigatti, Staiano et al., 2013, Menigatti, Truninger et al., 2009, Suzuki, Watkins et al., 2004). Furthermore, differentially methylated regions of the genome far outnumber gene mutations in precancerous colon tumors, and they tend to be found across most lesions (unlike mutations, which are likely to be tumor subset-specific (Dehghanizadeh, Khoddami et al., 2018, Lin, Raju et al., 2017, Nikolaev, Sotiriou et al., 2012)).

These considerations prompted us to quantitatively characterize genome-wide DNA hypermethylation in a prospectively collected series of SSA/Ps, each with a paired sample of normal mucosa, and to see how the epigenetic phenotype of SSA/Ps compares with those of cADNs and of CIMP colon cancers themselves. Our primary aim was to identify strong DNA methylation-based biomarker candidates with the potential for improving the identification and differential diagnosis of precancerous colorectal lesions. Our secondary aim was to determine the extent to which changes in the colon-precursor methylome are reflected in their gene expression profiles. To this end, we subjected the same tissue series to RNA-sequencing analysis, which also allowed us to identify numerous lesion-specific transcriptional dysregulations with potential for development as histopathologic biomarkers.

## Materials and Methods

### Analyses of prospectively collected tissues

#### Tissues

The fresh tissues used to study DNA methylation and gene expression were prospectively collected during colonoscopies performed between 2014 and 2017 at Cremona Hospital (Italy) or Zurich Triemli Hospital (Switzerland). The study was approved by both hospitals’ research ethics committees. Donors provided written consent to tissue collection, testing, and data publication. Samples were numerically coded to protect donors’ rights to confidentiality and privacy. Precancerous lesions were collected from the cecum or ascending colon during colonoscopies that were negative for cancers (**Table 1**). Each lesion was accompanied by a control sample of normal mucosa located >2 cm from the tumor (essential if baseline differences between tissue donors are to be accounted for). Lesions analyzed had: 1) maximum diameters of ≥ 10 mm (to ensure that sufficient tissue was left for the histological examination) and 2) Paris class Is or IIa morphologic features (i.e., sessile polyps and nonpolypoid lesions that were slightly elevated above the surrounding mucosa) (Endoscopic Classification Review Group, 2005) to increase the likelihood of including SSA/Ps. Tissue samples were placed in tubes filled with AllProtect Tissue Reagent (Qiagen, Hilden, Germany), held at 4°C overnight, and stored at -80°C prior to simultaneous DNA/RNA extraction with Qiagen’s AllPrep Mini Kit. All study tissues were histologically classified according to WHO criteria (Bosman et al., 2010) by an expert gastrointestinal pathologist at the hospital furnishing the lesion.

**Table 1:** Characteristics of the prospectively collected precancerous lesions.

| Patient number | Sex | Age | Colon segment involved | Maximum lesion diameter | Paris classification # | Pit pattern classification † | Microscopic appearance | Dysplasia Δ | No. of lesions at study colonoscopy § | *BRAF* | *KRAS* |
| --- | --- | --- | --- | --- | --- | --- | --- | --- | --- | --- | --- |
| S1 | F | 88 | C | 15 | 0-IIa | III<sub>s</sub> | SSA/P | none | 2 | c.1799T>A (V600E) | WT |
| S2 | F | 68 | A | 12 | 0-Is | III<sub>s</sub>, III<sub>L</sub> | SSA/P | none | 1 | c.1799T>A (V600E) | WT |
| S3 | F | 61 | A | 20 | 0-IIa | II | SSA/P | dysplasia | 4 | c.1799T>A (V600E) | WT |
| S4 | F | 71 | A | 15 | 0-IIa+II<sub>c</sub> | IV, V | SSA/P | dysplasia | 1 | c.1799T>A (V600E) | WT |
| S5 | M | 52 | C | 30 | 0-IIa | IV | SSA/P | none | 1 | c.1799T>A (V600E) | WT |
| S6 | F | 76 | C | 25 | 0-IIa | IV | SSA/P * | dysplasia | 2 | c.1799T>A (V600E) | WT |
| S7 | F | 59 | A | 14 | 0-IIa | nr | SSA/P | dysplasia | 2 | c.1799T>A (V600E) | WT |
| S8 | M | 63 | A | 14 | 0-Is | II | SSA/P | none | 6 | c.1799T>A (V600E) | WT |
| S9 | F | 69 | A | 20 | 0-IIa | II | SSA/P | none | 1 | c.1799T>A (V600E) | WT |
| S10 | F | 45 | A | 25 | 0-Is | Is | SSA/P | none | 1 | c.1799T>A (V600E) | WT |
| S11 | M | 57 | C | 14 | 0-IIa | III<sub>s</sub>, III<sub>L</sub> | SSA/P | none | 2 | WT | c.37G>C (G13R) |
| S12 | M | 49 | A | 10 | 0-IIa | nr | SSA/P | none | 3 | c.1799T>A (V600E) | WT |
| S13 | M | 63 | A | 15 | 0-Is | nr | SSA/P | none | 3 | c.1799T>A (V600E) | WT |
| S14<sup>o</sup> | F | 65 | A | 15 | 0-IIa | III<sub>s</sub> | SSA/P | none | 14 | c.1799T>A (V600E) | WT |
| S15 | F | 36 | A | 15 | 0-IIa | nr | SSA/P | none | 2 | c.1799T>A (V600E) | WT |
| S16 | M | 55 | C | 25 | 0-IIa+Is | III<sub>L</sub>, IV | SSA/P * | dysplasia | 0 | WT | c.35G>A (G12D) |
| S17 | F | 53 | C | 15 | 0-IIa | III<sub>s</sub> | SSA/P | none | 1 | c.1799T>A (V600E) | WT |
| A1 | M | 58 | A | 25 | 0-IIa | IV | TA | LGD | 1 | WT | c.38G>T (G13V) |
| A2 | M | 68 | A | 35 | 0-lps | IV | TVA | LGD | 1 | WT | c.37G>T (G13C) |
| A3 | F | 78 | A | 15 | 0-Is | IV | TA | LGD | 1 | WT | WT |
| A4 | F | 64 | A | 30 | 0-IIa+Is | IV | TA | LGD | 1 | WT | c.34G>T (G12C) |
| A5 | F | 59 | A | 25 | 0-IIa | III<sub>L</sub> | TA | LGD | 1 | WT | WT |
| A6 | F | 73 | A | 10 | 0-Is | III<sub>s</sub> | TA | LGD | 12 | WT | WT |
| A7 | F | 64 | C | 20 | 0-Is | III<sub>L</sub> | TA | LGD | 1 | WT | WT |
| A8 | M | 70 | A | 10 | 0-IIa | II, III<sub>s</sub> | TA | LGD | 3 | WT | WT |
| A9 | F | 73 | A | 30 | 0-IIa | III<sub>L</sub> | TA | LGD | 1 | WT | c.35G>T (G12V) |
| A10 | M | 77 | C | 40 | 0-Is | III<sub>L</sub>, IV | TA | LGD | 9 | WT | WT |
| A11 | F | 84 | A | 40 | 0-IIa | III<sub>L</sub>, IV | TA | LGD | 5 | WT | c.35G>T (G12V) |
| A12 | M | 61 | A | 25 | 0-lps | III<sub>L</sub>, IV | TVA | LGD | 1 | WT | WT |
| A13 | M | 83 | A | 18 | 0-Is | III<sub>L</sub> | TA | LGD | 1 | WT | WT |
| A14 | F | 78 | A | 15 | 0-Is | III<sub>s</sub> | TA | LGD | 2 | WT | WT |
| A15 | M | 75 | C | 20 | 0-IIa+Is | IV | TA | LGD | 1 | WT | c.37G>T (G13C) |

Abbreviations: M, male; F, female; C, cecum; A, ascending colon; TA, tubular adenoma; TVA, tubulovillous adenoma; SSA/P, sessile serrated adenoma/polyp; LGD, low-grade dysplasia; nr: not reported.

#### BRAF and KRAS mutation analysis

All samples were subjected to standard Sanger sequencing (Supplementary Experimental Procedures) to identify mutations in *BRAF* exon 15 (site of the mutational hotspot that gives rise to the V600E variant) and *KRAS* exon 2 (codons 12 and 13, which are frequently mutated).

#### Genome-wide bisulfite DNA sequencing and RNA sequencing

For methylation analysis, sequencing libraries were prepared using 1 microgram of DNA per sample, according to the Roche SeqCapEpi CpGiant protocol (Roche, Rotkreutz, Switzerland) (Supplementary Experimental Procedures), and bisulfite-converted prior to capture with a pre-designed probe pool (Roche-NimbleGen, Madison, WI). This target-enrichment procedure allowed us to interrogate 80.5Mb of the genome containing ˜2.7 × 10^6^ CpG sites, with thorough coverage of intra- and intergenic CpG islands and those in gene promoters, and more limited coverage of CpG shores and CpG nucleotides in enhancers and gene bodies. Captured libraries were sequenced on an Illumina 2500 system (Illumina, San Diego, CA) (125-bp paired-end reads).

RNA sequencing was restricted to samples with total RNA integrity numbers exceeding 6.5. Poly-A RNA was isolated from 100 ng of total RNA. PCR-amplified cDNA sequencing libraries were prepared according to the Illumina TruSeq Stranded mRNA Library preparation protocol and sequenced on an Illumina HiSeq 4000 system (150-bp paired-end reads).

#### Analysis of methylome and transcriptome data

Bisulfite- and RNA-sequencing reads were aligned to the GRCh37/hg19 human reference genome using Bismark/Bowtie 2 (Krueger & Andrews, 2011) and Salmon (Patro, Duggal et al., 2017), respectively. Quality-control analysis excluded one SSA/P (S14 in Table 1) from the methylome analysis because the probe pool’s target capture specificity in its matched normal mucosa sample was low. Differentially methylated cytosines (DMCs) (i.e. those displaying hyper- or hypo- methylation in tumor tissues vs. matched normal controls) were identified using the *BiSeq* R-package *(Hebestreit, Dugas et al., 2013).* Differentially methylated regions (DMRs) were defined as clusters of adjacent DMCs that were consistently hyper- or hypomethylated. (Data pre-processing, quality control, and *BiSeq* analysis are described in Supplementary Experimental Procedures.) Genes displaying significant differential expression in precancerous lesions (vs matched samples of normal mucosa) were identified using the *edgeR* R-package (Robinson, McCarthy et al., 2010) (details in Supplementary Experimental Procedures).

Raw methylome and transcriptome data are deposited in *ArrayExpress* (accession numbers: E-MTAB-6952; E-MTAB-6951; E-MTAB-6949).

### FFPE tissue studies

#### Tissues

Fresh-tissue findings on DNA methylation were verified in formalin-fixed, paraffin-embedded (FFPE) colon tissues (precancerous and normal mucosa) collected proximal to the splenic flexure. The tissues were obtained from the Zurich University Hospital Pathology Archives with local ethics committee approval and written donor consent (Supplementary Table 1). DNA was extracted from samples using the truXTRAC kit (Covaris, Woburn, MA) (Supplementary Experimental Procedures). FFPE samples were histologically classified and genotyped for *BRAF* and *KRAS,* as described above for the prospectively collected tissue series.

#### Bisulfite pyrosequencing

DNA methylation patterns at 6 selected DMRs were verified in the independent FFPE sample series with bisulfite pyrosequencing. DNA (500 ng per sample) was treated with sodium bisulfite using the Zymo Research DNA Methylation Lightning kit (Zymo Research, Irvine, CA). The bisulfite-converted DNA (20 ng) was amplified using biotinylated primers, and PCR products were pyrosequenced on a PyroMark Q24 Autoprep system (Qiagen) (Supplementary Experimental Procedures; Supplementary Table 2). The accuracy of the 6-marker panel in distinguishing the two types of precancerous lesions was calculated with a support vector machine model (Supplementary Experimental Procedures).

#### In-situ hybridization and immunohistochemistry

*In situ*-hybridization studies of representative lesions were performed with the RNAscope 2.0 assay (Advanced Cell Diagnostics, Hayward, CA), which uses multiple probes for each RNA and branched DNA molecules to amplify signals (Supplementary Experimental Procedures). MLH1 immunohistochemistry was performed on all lesions, as previously described (Truninger, Menigatti et al., 2005).

## Results

A total of 64 fresh colon tissue samples—17 histologically classified as SSA/Ps and 15 considered cADNs (Table 1), each with matched samples of normal mucosa—were subjected to both genome-wide bisulfite sequencing and RNA sequencing. The BRAFV600E mutation was found in 15 (88%) of the SSA/Ps and none of the cADNs. *KRAS* mutations were present in *BRAF*-wildtype SSA/Ps and in 6 (40%) of the 15 cADNs. As expected for early-stage serrated tumors, none of the SSA/Ps exhibited immunohistochemical evidence of *MLH1* silencing.

### Proto-CIMP SSA/Ps

**Figure 1A** shows the results of unsupervised, multidimensional scaling (MDS) analysis of the DNA methylomes of the fresh-tissue series, as determined by genome-wide bisulfite sequencing. For comparison purposes, we used the same method to re-analyze DNA samples from 6 proximal-colon cancers (and matched normal mucosa samples) collected in a previous study (di Pietro, Sabates Bellver et al., 2005) (Supplementary Table 3). The methylation data neatly segregated neoplastic tissues from their paired normal controls (dimension 1) and SSA/P from cADN samples (dimension 2). As for the cancers, CIMP(+) (corresponding to the CIMP-high subgroup of ref. (Hinoue et al., 2012)) and CIMP(-) (i.e., non-CIMP) tumors clustered with the SSA/Ps and cADNs, respectively.

**Figure 1:**
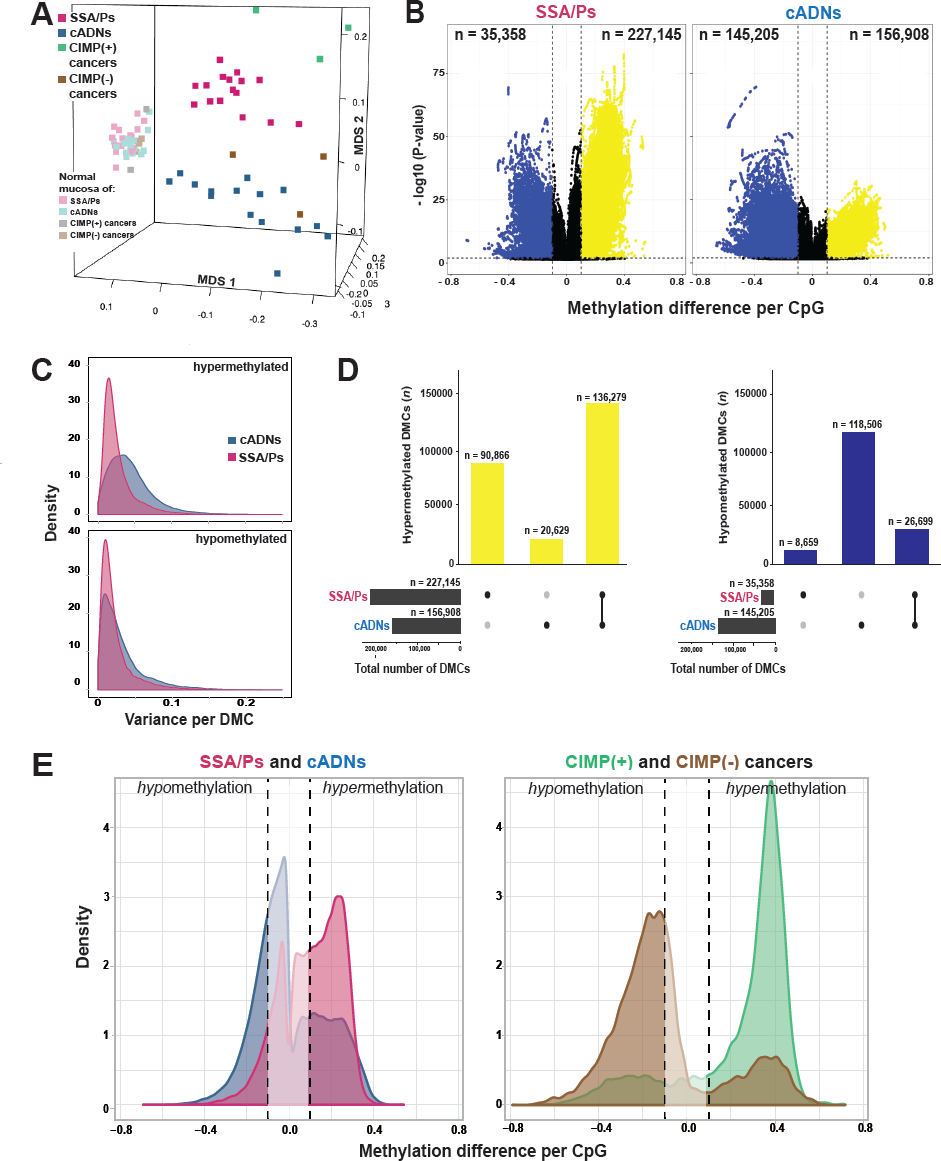
Differentially methylated cytosines (DMCs) in precancerous and cancerous colon lesions. **A.**MDS plot (Supplementary Experimental Procedures) of DNA methylation levels in SSA/Ps (n=16); cADNs (n=15); CIMP(+) cancers (n=3); CIMP(-) cancers (n=3); and matched samples of normal mucosa for each lesion. **B.** Volcano plots showing the magnitude (x axis) and statistical significance (y axis) of the differential methylation observed at DMCs identified in SSA/Ps and cADNs. X-axis: The magnitude of differential methylation was calculated as the M:T ratio (no. methylated reads / total no. reads) for the tumor sample minus M:T ratio for matched normal-tissue control. Black dots: DMCs with absolute methylation differences of <0.1 and P-values >0.01. Yellow and blue dots: highly significant (P-value <0.01) DMCs (hypermethylated and hypomethylated, respectively). **C.** Density plot showing variance at the hypermethylated (top) and hypomethylated (bottom) DMCs (yellow and blue dots of panel B, respectively). **D**. UpSet plots showing hypermethylated (left) and hypomethylated (right) DMC sets in SSA/Ps and cADNs and their overlaps. Exact numbers of lesion-specific (•) and shared (•-•) DMCs appear above the bars. **E.** Overlaid density plots showing the distributions of hypo- and hypermethylated DMCs in SSA/Ps and cADNs (left), and CIMP(+) and CIMP(-) cancers (right). X-axis as described in panel B.

Cytosines displaying differential methylation in the precancerous lesions relative to their counterparts in matched normal tissues (DMCs) abounded in both SSA/Ps (*n*=262,503) and cADNs (*n*=302,113). *Hypo*methylated DMCs were much more frequent in cADNs, while hypermethylated DMCs were more common in SSA/Ps (**Figures 1B** and **D**). The lower *P*-values for the hypermethylated DMCs in SSA/Ps probably reflect the more uniform hypermethylation levels seen across these lesions (**Figure 1C**). Forty percent of the hypermethylated DMCs in SSA/Ps were unique to these precancers, while 87% of those in cADNs were also present in SSA/Ps (Figure 1D). Consistent with findings shown in Figure 1A, the CIMP(+) cancers also displayed more hypermethylation and less hypomethylation than their CIMP(-) counterparts, although both types of differential methylation were more striking in the cancer methylomes than in those of the precursors (**Figure 1E**), a pattern that reflects the increasing divergence of the lesions’ methylation profiles as they move progressively along the two different tumorigenic pathways.

As for DMRs, those *hyper*methylated were, as expected, more numerous in SSA/Ps, while *hypo*methylated DMRs were much more common in cADNs (**Figure 2A**). Most hypermethylated DMRs were detected in both types of precancers, but a substantial number were SSA/P-specific (**Figure 2B**). By contrast, hypomethylated regions tended to be cADN-specific (Figure 2B). Hypomethylation was also more likely to be found outside CpG-rich areas and within introns and intergenic regions (**Figure 2C).** In both cancer precursor classes, hypermethylated DMRs were located prevalently in CpG islands / shores and gene promoters.

**Figure 2:**
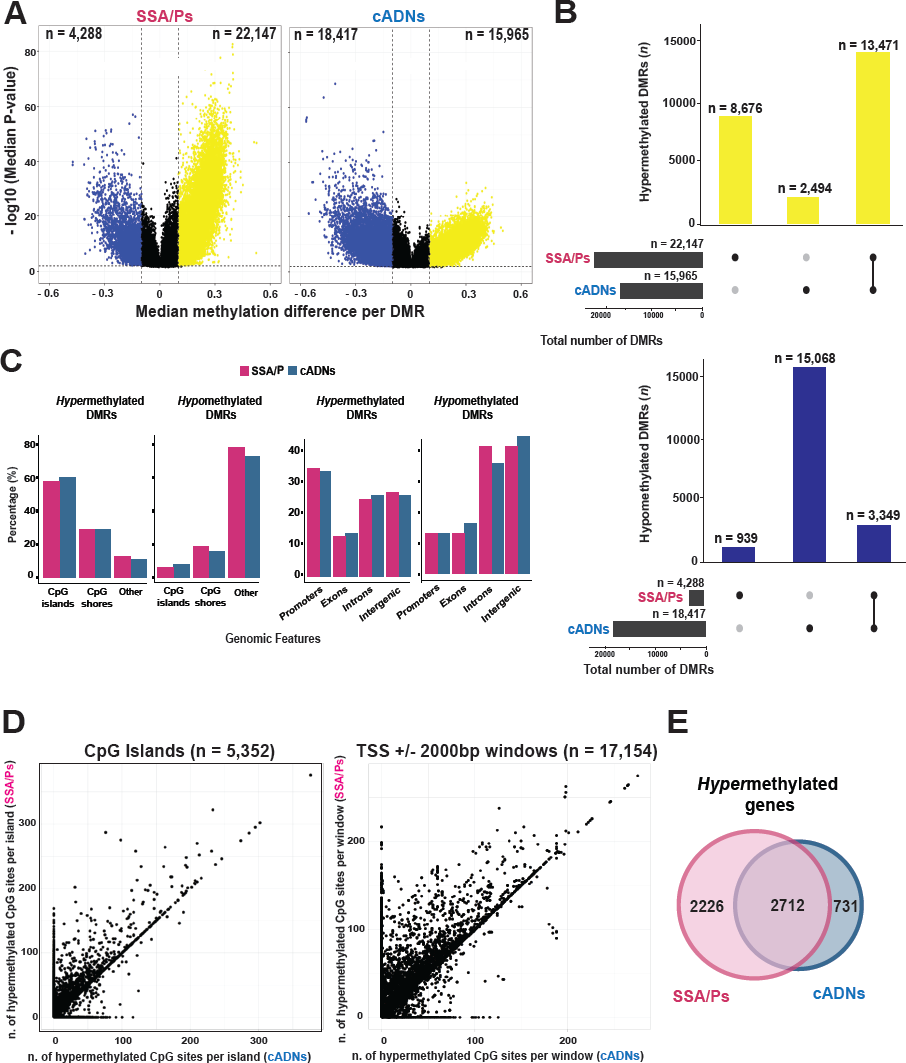
Differentially methylated *regions* (DMRs) in precursor lesions and proto-CIMP in SSA/Ps. **A.** Volcano plots showing the median differential methylation (*x-axis*) and its statistical significance (*y-axis*) for DMRs found in SSA/Ps and cADNs. (See Figure 1B legend for cutoffs and color codes) **B.** UpSet plots showing the lesion-specificity of hypermethylated and hypomethylated DMRs found in precancerous tumors. Exact numbers of lesion-specific (•) and shared (•-•) DMRs appear above the bars. **C.** Genomic location of DMRs in SSA/Ps vs cADNs. **D.** Scatterplots showing the extent of methylation at regions hypermethylated in both SSA/Ps and cADNs, and located within CpG islands (dots in the left panel) or 4-kb peri-TSS windows, i.e., TSS plus 2000-bp upstream and downstream flanking regions (dots in the right panel). In addition, CpG islands or peri-TSS windows with at least one hypermethylated CpG in either SSA/Ps or cADNs are included in the two graphics (i.e., dots along the *y* or *x* axis, respectively). **E.** Venn diagrams showing overlap between sets of genes in SSA/Ps and cADNs whose peri-TSS window contained at least one hypermethylated DMR.

Next, we compared the 5352 CpG island-associated DMRs and 17,154 transcription start site (TSS)-associated DMRs (peri-TSS DMRs) that were hypermethylated in both classes of precursors. Interestingly, those present in SSA/Ps contained appreciably higher numbers of hypermethylated CpGs (**Figure 2D** and the example in Supplementary Figure 1). Furthermore, some SSA/P-specific DMRs displaying relatively mild hypermethylation seemed to correspond to longer, more intensely methylated regions seen in CIMP(+) colon cancers (e.g., the large CpG island shared by the bidirectional promoters of the *MLH1* and *EPM2AIP1* genes). As shown in Supplementary Figure 2A, aberrant methylation of this island involves both the *EPM2AIP1* and *MLH1* promoters in CIMP(+) cancers, where the *MLH1* expression loss triggers MMR deficiency. In SSA/Ps, the hypermethylation is instead confined to the *EPM2AIP1* promoter and associated with a decrease in the expression of this gene (Supplementary Figures 2B and C). While methylation at this locus has been investigated in cell lines (Lin, Jeong et al., 2007), tissue-based data are lacking on the specific region where the serrated tumor-associated hypermethylation appears to begin.

As shown in **Figure 2E,** genes whose peri-TSS window included at least one hypermethylated DMR were appreciably more common in SSA/Ps than cADNs. (Affiliation of DMRs to genes is described in Supplementary Experimental Procedures; the genes themselves are listed in Supplementary Table 4.)

Collectively, the data presented above indicate that distinctive methylome features of CIMP(+) colon cancers are readily detectable in their putative precursors, SSA/Ps (albeit in less marked forms). Therefore, this early-stage hypermethylator phenotype will be referred to hereafter as *proto-CIMP.*

### Targeted verification of DMR-based biomarker candidates for identifying colon-cancer precursors

To explore their potential as DNA-based diagnostic markers, we performed targeted verification studies on 6 of the hypermethylated DMRs listed in Supplementary Table 4 (3 that were SSA/P-specific and 3 others shared by SSA/Ps and cADNs). The selection was subjective and based on visual inspection of the methylation patterns across all samples in the Integrative Genomics Viewer (IGV). Factors considered included pattern variation across tumor samples and baseline methylation levels in paired normal mucosa samples. The hypermethylation patterns in our freshly-collected tissue samples of the 6 biomarker candidates and those of other DMRs currently utilized as colon-cancer markers in clinical research (Hinoue et al., 2012, Weisenberger et al., 2006) are shown in Supplementary Figure 3.

Bisulfite pyrosequencing assays were developed to verify our biomarker candidates in the independent series of FFPE samples (Supplementary Table 1). The results (**Figure 3A** and **B**) fully confirmed genome-wide bisulfite DNA sequencing data on single-CpG methylation levels within each DMR candidate. The 6-marker panel distinguished precancerous lesions from normal mucosa and SSA/Ps from cADNs (**Figure 3C**) with 96.7% accuracy (Supplementary Figure 4A). Preliminary data on a small number of FFPE samples of TSAs and colon cancers analyzed with this panel (Supplementary Figure 4B) suggest that the TSA methylome more closely resembles that of cADNs than that of SSA/Ps and confirm findings shown in Figure 1A, whereby colon cancer clustering with SSA/Ps or cADNs appears to be CIMP status-dependent.

**Figure 3:**
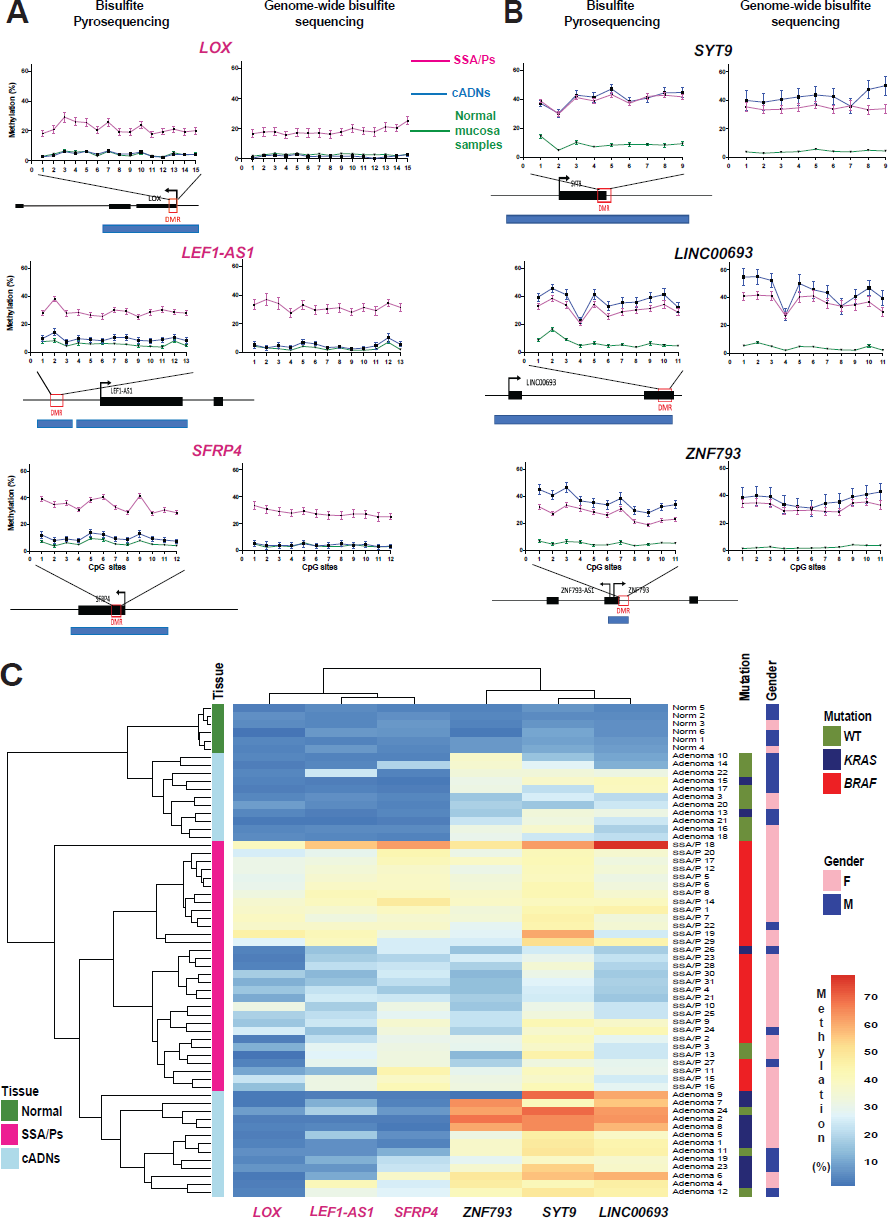
Bisulfite pyrosequencing verification of 6 hypermethylated DMRs. Methylation levels at each CpG site of **(A)** 3 SSA/P-specific DMRs and **(B)** 3 DMRs shared between SSA/Ps and cADNs. Verification assay results in 61 FFPE colon tissue samples (left) are compared with those of genome-wide bisulfite sequencing of the fresh tissue series (right). DMR-containing loci (red box) and CpG islands (blue horizontal bars) are schematically represented below graph pairs. **C.** Hierarchical clustering heatmap of the 61 FFPE tissue samples based on the mean methylation level across all CpGs in each DMR.

### The SSA/P and cADN transcriptomes

To explore how the methylome profiles reported above are reflected in the transcriptomes of precancerous colon lesions, we performed RNA sequencing-based gene expression analyses on the 64 fresh-tissue samples used for methylation profiling. The results readily segregated most precancers into SSA/Ps and cADNs (**Figure 4A** and **B**). MetaCore enrichment analysis of the most strikingly dysregulated genes in the precancers (**Figure 4C** and Supplementary Table 5) identified pathways involved in “Cell adhesion_extracellular remodelling” and “Development_WNT signaling” as the ones most markedly altered in SSA/Ps and cADNs, respectively.

**Figure 4:**
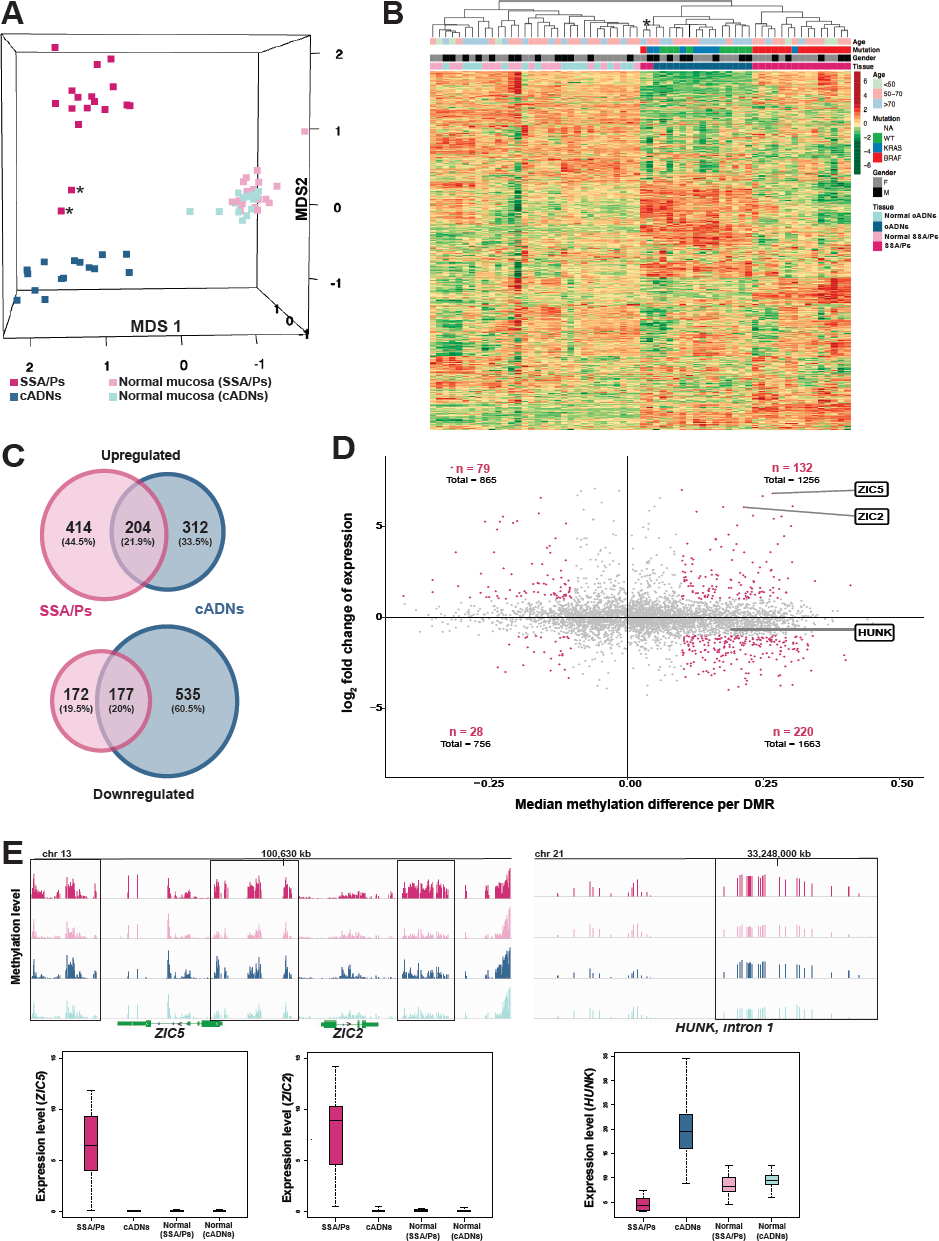
Analysis of the SSA/P and cADN transcriptomes in light of the lesions’ methylome profiles. **(A)** MDS plot and **(B)** hierarchical clustering heatmap of the 17 SSA/Ps and 15 cADNs and their normal mucosal samples. Asterisks indicate the two samples with intermediate profiles (for details, see Table 1). Hierarchical clustering in B was based on expression levels of the 10,000 genes with most highly variable expression. **C.** Venn diagrams showing the precursor-lesion specificity of the most dysregulated genes (P-value <10^-10^ and log2 fold change >1). **D.** Scatter plots showing the variable relation between the magnitude of differential DNA methylation (x axis) and the expression (y axis) for SSA/Ps (vs matched normal mucosa samples). Genes with at least one DMR (P-value <0.05) in their peri-TSS windows and dysregulated expression (P-value <0.05) are shown. Red dots: subsets of genes with a median methylation differences of ≥ 0.1 and log_2_ fold change in expression of >1. *ZIC2, ZIC5,* and *HUNK* are highlighted to illustrate the heterogeneous relation between DNA methylation and gene expression. **E.** Methylation levels (top) and gene expression (bottom) data for the *ZIC2*, *ZIC5,* and *HUNK* loci in SSA/Ps (pink), cADNs (blue), and corresponding samples of normal mucosa (light-pink and light blue, respectively). Areas outlined in black are those displaying differential methylation (see details in Supplementary Figures 5 and 6).

As shown in Figure 4C, the major gene expression alterations in cADNs were usually downregulations, whereas increased expression was more common in SSA/Ps. The predominance of upregulated transcription in lesions displaying proto-CIMP illustrates the emerging complexities of DNA methylation’s relation to transcription. Although hypermethylation of promoter CpG islands is frequently cited as a silencing epigenetic mark (Jones, 2012), the relation between peri-TSS methylation and gene expression levels in our samples varied widely. For example, as shown in **Figure 4D,** substantial hypermethylation was associated with significantly upregulated expression of *ZIC2* and *ZIC5* but mildly reduced expression of *HUNK*. Inverse (hypermethylation with downregulation, hypomethylation with upregulation) and positive (hypermethylation with upregulation, hypomethylation with downregulation) relations were encountered with similar frequencies. Furthermore, significantly dysregulated expression was also observed for many genes which—unlike those represented in Figure 4D—had no evidence of aberrant promoter methylation. **Figure 4E** (and Supplementary Figures 5 and 6) provide more details on the relations between gene expression and DNA methylation at the *ZIC2, ZIC5,* and *HUNK* loci. *In situ* hybridization-based tissue staining patterns for *ZIC2* and *HUNK* mRNAs in precancerous lesions are shown in **Figure 5**.

**Figure 5:**
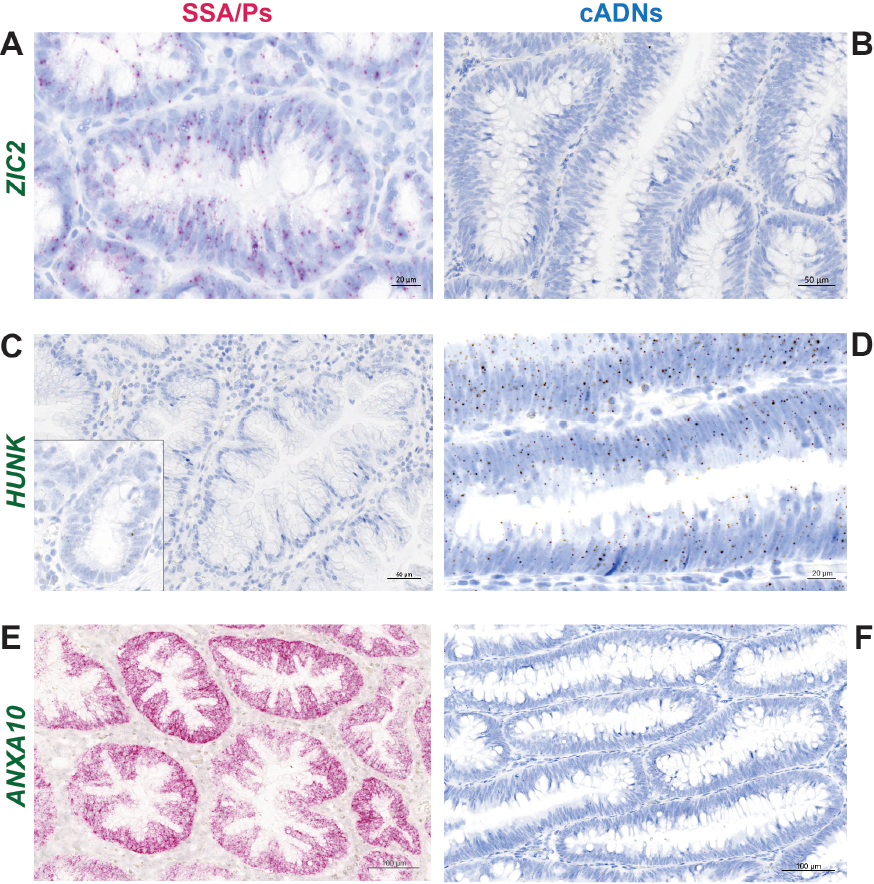
*In situ* hybridization analysis of *ZIC2*, *HUNK,* and *ANXA10* mRNAs in SSA/Ps and cADNs. *ZIC2* expression (red punctate labeling) is **(A)** present in SSA/Ps and **(B)** absent in cADNs. *HUNK* (brown punctate labeling) is **(C)** almost absent in SSA/Ps (very low-level expression was present at the bottom of serrated crypts, see inset) and **(D)** expressed in cADNs. Control staining for *ANXA10,* a known SSA/P-specific marker (Delker et al., 2014, Kim et al., 2015, Sajanti et al., 2015, Sakamoto et al., 2017) was (**E**) strongly positive in SSA/Ps (dense red punctate labeling) and (**F**) absent in cADNs.

## Discussion

Previous studies have analyzed DNA methylation alterations in colon *cancers* with an eye to their exploitation as diagnostic markers. In 2006, Weisenberger et al. analyzed methylation at 195 genetic loci and identified a panel of 5 DMRs that discriminated between CIMP and non-CIMP colon cancers (Weisenberger et al., 2006). This panel was refined by the same group based on the analysis of ˜27,000 CpG sites (Hinoue et al., 2012). Although several groups have also used these panels (and other previously-identified sets of cancer-associated methylation markers (Toyota et al., 1999)) to classify *precancerous* colon lesions (Burnett-Hartman, Newcomb et al., 2013, Dhir, Yachida et al., 2011, Fernando, Miranda et al., 2014, Kambara et al., 2004, Kim, Kakar et al., 2008, Maeda, Suzuki et al., 2011, O’Brien et al., 2006, Park, Rashid et al., 2003, Vaughn, Wilson et al., 2010, Worthley, Whitehall et al., 2010, Yamamoto, Suzuki et al., 2012), their diagnostic potential in this setting is uncertain. While some regions might be aberrantly methylated from the outset of colon tumorigenesis, others could conceivably be present in cancers but absent or poorly developed in early-stage colon tumors (e.g., that affecting the *MLH1* promoter).

Dehghanizadeh et al. recently attempted to explore this uncharted territory using a microarray-based assay to assess methylation at ˜450,000 CpG sites (Dehghanizadeh et al., 2018). However, the series of precancerous lesions they studied included only 5 SSA/Ps (archival samples: fresh or FFPE) and 3 cADNs (all from patients with familial polyposis), and, with the exception of one of the SSA/Ps, none of the precancerous lesions were accompanied by a paired sample of normal mucosa. Our methylation analysis included a total of 64 prospectively collected, fresh-tissue samples representing the two major classes of sporadic precancerous colon lesions, SSA/Ps and cADNs, and patient-matched samples of non-neoplastic tissues. The results show that DMR profiles can distinguish colon cancer precursors from normal colon tissues, and they can also differentiate between the SSA/Ps and cADNs. Hypermethylation within gene regulatory regions—a common feature of tumorigenesis in general—was encountered frequently in both types of precancers. However, in SSA/Ps it is appreciably more pervasive, and it shares many features (albeit in milder forms) with the hypermethylation typical of the CIMP(+) colon cancers believed to develop from SSA/Ps—hence our referral to the SSA/P phenotype as *proto-CIMP*. Global hypomethylation of the genome is also a common feature of tumorigenesis, and it was clearly evident in both cADNs and CIMP(-) colon cancers. It has also been reported in a single SSA/P investigated with whole-genome bisulfite sequencing (Dehghanizadeh et al., 2018). However, our data indicate that SSA/Ps and CIMP(+) cancers are generally spared from the wave of demethylation that occurs during conventional colorectal tumorigenesis (at least within the 80.5 Mb area of the genome we investigated). This epigenetic phenotype might thus be a novel feature of CIMP tumorigenesis. As such, it clearly deserves further investigation with a whole-genome study of methylation in several precursor lesions of different types.

The six markers verified in this study (*LOX*, *LEF1-AS1*, *SFRP4*, *ZNF793*, *SYT9,* and *LINC00693*) displayed high accuracy (97%) in distinguishing the colon precancers from normal mucosa and also in differentiating between the two main classes of precancerous tumors. However, the long list of DMRs we identified (Supplementary Table 4) should be scrupulously mined to find additional markers, which can be used to develop a stool-DNA test that excels in both sensitivity (i.e., detection of precancerous as well as cancerous lesions) and specificity (i.e., differentiation of SSA/Ps / CIMP(+) cancers and cADNs / CIMP(-) cancers). Indeed, the vast majority of DMRs we identified in the precancers, including the 6 markers, appear to undergo little or no negative selection during transformation and therefore were also present in colon cancers. Recently, a stool-DNA test, whose targets include aberrant *BMP3* and *NDRG4* methylation, has produced promising results in the pre-colonoscopy detection of cancers and advanced precancerous lesions of the colon (including advanced SSA/Ps) (Imperiale, Ransohoff et al., 2014). As shown in Supplementary Figures 3Q and R, these markers, which were also in our list, performed well in the identification of our CIMP(+) and proto-CIMP tumors.

The results of our RNA sequencing studies confirm the increasing body of evidence highlighting the complex relation between DNA methylation and gene expression (Jones, 2012). Emphasis in the literature has frequently been placed on the association between gene silencing and hypermethylation of promoter CpG islands, since the latter is an established mechanism underlying cancer-related inactivation of tumor suppressor genes (e.g., *MLH1* in colon tumorigenesis) (Herman, Umar et al., 1998, Lin et al., 2007, Truninger et al., 2005). Analysis of our data failed to reveal any genes with established tumor-suppressor functions among those with downregulated expression and peri-TSS hypermethylation in precancerous lesions. *HUNK*, for example, encodes a protein kinase that appears to negatively regulate normal intestinal cell proliferation (Reed, Korobko et al., 2015), but it also seems to promote mammary tumorigenesis (Yeh, Belka et al., 2013). As for its methylation pattern, a substantial portion of *HUNK*’s peri-TSS window was hypermethylated in the CIMP(+) cancers. In contrast, the methylation observed in SSA/Ps was reduced in level and confined to a shorter stretch of CpG sites. (The latter feature recalls the aberrant methylation involving the CpG island shared by the *EPM2AIP1* and *MLH1* promoters, which affected the MMR gene promoter only in the more advanced stages of tumorigenesis.) The methylation present in the SSA/Ps was nonetheless associated with mildly reduced *HUNK* expression (Figure 4E and Supplementary Figure 5A). Interestingly, *HUNK* methylation (even milder than that seen in SSA/Ps) was also identified in some cADNs, where it was associated with *upregulated* expression of the gene (Supplementary Figure 5B).

Consistent with previous reports (Bergman & Cedar, 2013, Hinoue et al., 2012), the CpG island hypermethylation events we detected mainly affected the promoter regions of genes displaying little or no expression in precancerous and normal mucosa. Some of these genes are known to be marked during embryonic development by polycomb repressive complex 2 (PRC2), which mediates H3K27 trimethylation, a hallmark of gene silencing. During tumorigenesis, these loci display increases in DNA methylation together with decreases in H3K27me3. The mechanisms responsible for these changes are incompletely understood (Manzo, Wirz et al., 2017, Mohn, Weber et al., 2008, Reddington, Pennings et al., 2013, Schlesinger, Straussman et al., 2007, Widschwendter, Fiegl et al., 2007), but they are believed by some to tighten repression of the target gene’s expression, rendering it more difficult to induce (Easwaran, Johnstone et al., 2012).

Notably, however, the markedly dysregulated gene expression in our SSA/Ps tended to involve upregulation rather than downregulation (Figure 4C), despite the lesions’ proto-CIMP. Indeed, several upregulated genes exhibited hypermethylation in their proximal regulatory regions (Figure 4D). For example, the *ZIC2* / *ZIC5* locus (Figure 4 and details in Supplementary Figure 6) was methylated strongly and extensively in SSA/Ps—and less so in cADNs—and yet, the expression of these two genes was re-activated only in the SSA/Ps (after PRC2 repression during embryonic development). Additional work is needed to unravel the ties between DNA methylation and gene expression at this locus (and others) in SSA/Ps and cADNs. The fairly small differences in the two lesions’ hypermethylation patterns might well be associated with more marked differences at the chromatin level.

Gene expression is also modulated by *cis* epigenetic signals other than DNA methylation, including myriad histone modifications, and by long-range regulatory signals, such as those classically involving enhancers via CTCF-mediated chromatin loop formation (Ong & Corces, 2014). And, naturally, differential transcription-factor expression in the two types of lesions will be reflected in these proteins’ differential binding patterns in gene regulatory regions. The temporal coordination of these factors is highly complex and involves substantial backward regulation. For example, the binding affinity of many transcription factors—and CTCF—varies with the methylation status of their binding sites and, contrary to common belief, many factors preferentially bind methylated DNA (Flavahan, Drier et al., 2016, Wang, Maurano et al., 2012, Yan, Tang et al., 2016, Yin, Morgunova et al., 2017).Therefore, 5-methyl-cytosine nucleotides modulate protein-DNA interactions: the abnormal DNA methylation patterns in SSA/Ps and cADNs create alternative platforms for recruiting DNA- and chromatin-binding proteins. For example, the markedly reduced binding affinities of PRC2 and CTCF at hypermethylated DMRs causes these proteins to relocate to other genomic regions, giving rise to long-range repression of gene expression and changes in topologically associated genomic domains, respectively. Conversely, the re-expression of transcription factors, such as ZIC2 and ZIC5 (whose roles in tumorigenesis are completely unknown) specifically in SSA/Ps, would lead to additional long-range *trans* regulation of genes whose enhancers and promoters have binding sites for them.

Regardless of their functional relations with our DNA methylation data, the transcriptome differences we documented between SSA/Ps and cADNs can also be fruitfully exploited to develop tissue staining assays to differentiate these lesions with confidence. The extensive list of genes in Supplementary Table 5 represents a rich source of promising candidate markers that can be used for this purpose. For tissue staining, the recently developed *in situ* hybridization method we employed is straightforward and produces highly sensitive and specific results. It is a valuable alternative when specific antibodies are unavailable or do not perform well in immunohistochemistry, and excellent for tissue staining of non-coding RNAs.

Our list of potential expression-based SSA/P markers shows overlap with those identified by other groups (e.g., *ANXA10*, which is reportedly a highly specific marker of serrated-pathway colon tumors (Delker, McGettigan et al., 2014, Kim, Kim et al., 2015, Sajanti, Vayrynen et al., 2015, Sakamoto et al., 2017)). Indeed, our transcriptomic data appreciably extend previously published microarray findings (Caruso, Moore et al., 2009, De Sousa E Melo, Wang et al., 2013, Gonzalo, Lai et al., 2013) and are similar to RNA sequencing-based results from another group (Delker et al., 2014, Kanth, Bronner et al., 2016) (Supplementary Figure 7A). However, separate signatures are also needed for other types of serrated lesions and lesions typically found outside the proximal colon (hyperplastic polyps in particular, which are encountered very frequently). These areas are already being explored in several studies, (Kanth et al., 2016, Kim et al., 2015, Rahmatallah, Khaidakov et al., 2017) including the previously mentioned study based on RNA sequencing, which showed that SSA/Ps can be differentiated from hyperplastic polyps based on transcriptome profiles (Supplementary Figure 7B) (Kanth et al., 2016).

In conclusion, our study represents the most comprehensive genome-wide comparison of DNA methylation and gene expression in SSA/Ps and cADNs. The 6 verified markers in this study could be supplemented with other promising candidates from our DMR dataset and validated in future studies on a larger series of precancerous lesions. As with the gene expression, the sample set should be extended to involve all types of colonic tumors, including those from the distal colon. The findings from such studies will have clinical implications for the pre-colonoscopy stool-DNA-based detection— and identification—of early-stage colon tumors and for their more accurate molecular histological diagnosis.

### The Paper Explained

#### PROBLEM

Most colon cancers arise from conventional adenomas (cADNs), but those with the CpG island methylator phenotype (CIMP) appear to descend from sessile serrated adenomas/polyps (SSA/Ps). Compared with cADNs, SSA/Ps are more difficult to visualize and resect during colonoscopy, and their histological diagnosis is subject to more substantial inter-observer variability. To identify novel fecal and/or tissue-based biomarkers capable of improving the diagnosis and clinical management of SSA/Ps, we used high-throughput DNA methylation and gene expression analysis to investigate prospectively collected samples of SSA/Ps, cADNs, and paired specimens of normal mucosa.

#### RESULTS

Compared with matched samples of normal colon mucosa, SSA/Ps and cADNs both display markedly remodeled methylomes. SSA/Ps lack the widespread hypomethylation that prevails in cADNs, displaying instead a “proto-CIMP,” i.e., pervasive, extensive DNA hypermethylation resembling a milder version of the CIMP that characterizes their descendent cancers. A test panel of six selected hypermethylated regions (SSA/P-specific [n=3], present in SSA/Ps and cADNs [n=3]) distinguished precancers from normal mucosa and SSA/Ps from cADNs with high overall accuracy (96.7%). Surprisingly, proto-CIMP was associated with upregulated rather than silenced gene expression.

#### IMPACT

The strikingly different epigenetic landscapes of SSA/Ps and cADNs can be exploited to develop novel tissue-based and noninvasive biomarkers that could improve the detection and typing of precancerous colon lesions.

Author contributions
G.M. and H.R.P. designed research; H.R.P. and A.M.O. performed research; S.O. and H.R.P. analyzed data; F.C., H.H., M.S., S.V., L.A., and F.B. performed tissue sample and clinical data collection; G.T., A.W., and P.K. reviewed histology; H.R.P. and G.M. wrote the paper.

## Acknowledgments

We thank Martin Roszkowski, Vera van der Weijden, Hadi Gharibi, Anette Hunziker, Catharine Aquino, and Brigitta Tomlinson for their technical support, help, and productive discussions, and Marian Everett Kent for editing the manuscript.

## Abbreviations

cADNs: Conventional adenomas
CIMP: CpG island methylator phenotype
DMCs: Differentially methylated cytosines
DMRs: Differentially methylated regions
FFPE: Formalin-fixed, paraffin-embedded
IGV: Integrative genomics viewer
MDS: Multidimensional scaling
MMR: DNA-mismatch repair
PRC2: Polycomb repressive complex 2
SSA/Ps: Sessile serrated adenomas/polyps
TSAs: Traditional serrated adenomas
TSS: Transcription start site
WHO: World Health Organization

